# Closely related *Cryptococcus neoformans* strains possess differential virulence both in humans and the mouse inhalation model

**DOI:** 10.1101/524165

**Authors:** Liliane Mukaremera, Tami R. MacDonald, Judith N. Nielsen, Andrew Akampulira, Charlotte Schutz, Kabanda Taseera, Conrad Muzoora, Graeme Meintjes, David B. Meya, David R. Boulware, Kirsten Nielsen

**Author notes:** Corresponding Author: Kirsten Nielsen, PhD, Professor, Department of Microbiology and Immunology, University of Minnesota, 689 23^rd^ Ave SE, Minneapolis, MN 55455.

## Abstract

Cryptococcal meningitis (CM) causes high rates of HIV-related mortality, yet *Cryptococcus* factors influencing patient outcome are not well understood. Pathogen-specific traits, such as the strain genotype and degree of antigen shedding, are associated with clinical outcome but the underlying biology remains elusive. In this study, we examined factors determining disease outcome in HIV-infected cryptococcal meningitis patients infected with *C. neoformans* strains with the same multi-locus sequence type. Both patient mortality and survival were observed during infections with the same sequence type. Disease outcome did not correlate with underlying patient immune deficiencies. Patient mortality was associated with higher antigen levels, fungal burden in the CSF, and low CSF fungal clearance. Virulence of a subset of clinical strains with the same sequence type were analyzed using the mouse inhalation model of cryptococcosis. We showed a strong correlation between human and mouse mortality rates, demonstrating the mouse inhalation model recapitulates human infection. Similar to human infection, the ability to multiply *in vivo*, demonstrated by high fungal burden in the lung and brain tissues, was associated with mouse mortality. Mortality rate was not associated with single *C. neoformans* virulence factors *in vitro* or *in vivo*; rather, a trend in mortality rate correlated with a suite of traits. These observations show that genotype similarities between *C. neoformans* strains do not necessarily translate into similar virulence either in the mouse model or in human patients. In addition, our results show that *in vitro* assays do not fully reproduce *in vivo* conditions that influence *C. neoformans* virulence.

## Introduction

*Cryptococcus neoformans* is a fungal pathogen that causes disease mainly in immunocompromised patients such as individuals living with HIV/AIDS or receiving organ transplants. The availability of antiretroviral therapy has reduced AIDS-related mortality, however deaths due to cryptococcal meningitis (CM) have plateaued with *C. neoformans* still causing 15 *%* of all AIDS-related deaths globally (1, 2). Mortality rates due to *Cryptococcus* infection varies by region from 70% in low-income countries to 20-40% of all infected patients in high-income countries (2). Although differences in mortality rates between high- and low-income countries can be linked to sub-optimal antifungal treatments in low-income countries (3), variations in mortality rates between patient groups receiving similar treatments and residing in the same region of the world are observed (1, 4–6). Mortality is a measure influenced by many intrinsic host and pathogen factors (as well as host-pathogen interaction factors). Thus, we need better proxies to understand *C. neoformans* pathogenesis in patients and identify factors determining clinical outcome.

Although *C. neoformans* pathogenesis in the mouse model of cryptococcosis has been extensively studied, the correlation between disease characteristics in the human patient and clinical isolate infection in mice is unknown. The mouse model has been used to define *C. neoformans* virulence factors that allow the organism to be pathogenic, such as capsule, melanin, titan-cell formation, etc. The progression of disease in the inhalational mouse model - from initial inhalation into the lungs, to disseminated disease, and ultimately death due to central nervous system infection – is similar to human infection. Yet whether the mouse model accurately recapitulates differences in human disease, and whether the model can be used to identify subtle variations in virulence factors that impact the outcome of human disease remains unexplored. Having clinical information from patients, we investigated the association between disease parameters in patients compared to mice infected with the same *C. neoformans* isolate.

Previous studies showed that pathogen-specific characteristics, such as genotype or the degree of antigen shedding, influence immune responses to *C. neoformans* and the clinical outcome of infected patients (5, 7–9). Multi-locus sequence typing (MLST) of 7 genetic loci has been used previously to identify genetically similar strains (5–10). For example, studies of clinical isolates in both Uganda and Brazil showed higher patient mortality associated with sequence type 93 (ST93) strains (5, 10). ST93 strains had increases in type 2 cytokines in *ex vivo* cytokine release assays, suggesting that these strains may shift the Th1/Th2 immune balance (5). Similarly, studies from Vietnam have identified an association between ST5 and infections in non-HIV patients but the underlying biological differences remain unknown (11). To explore the association between genotype and clinical outcome, we examined individual patient differences in outcome and immune response in patients infected with the same ST type. We found human patients infected with *C. neoformans* strains with the same sequence type can have dramatically different clinical outcomes. This dichotomy in disease outcome was recapitulated in the mouse model of cryptococcosis, suggesting these differences in clinical outcome were due to strain-specific characteristics.

## Materials and Methods

### Ethical Statement

Animal experiments were done in accordance with the Animal Welfare Act, United States federal law, and NIH guidelines. Mice were handled in accordance with guidelines defined by the University of Minnesota Animal Care and Use Committee (IACUC) under protocol 1308-30852A.

The study population consisted of human immunodeficiency virus (HIV)-infected, antiretroviral therapy (ART)-naive individuals with a first episode of cryptococcal meningitis screened for the Cryptococcal Optimal ART Timing (COAT) trial (clinicaltrials.gov: NCT01075152) (12). Participants were enrolled from two countries; Uganda (Mulago Hospital in Kampala and Mbarara Hospital in Mbarara) and South Africa (GF Jooste Hospital in Cape Town) between November 2010 and April 2012. Written informed consent was obtained from all subjects or their proxy, and all data were de-identified. Institutional Review Board approvals were obtained from each participating site. A total of 106 patients that had culture-positive CSF for *Cryptococcus* and from which isolates were sequenced for genotypic analyses (5, 8) were included in this study.

### Strains and media

*Cryptococcus* clinical isolates were colony-purified from the cerebrospinal fluid (CSF) specimens from patients enrolled in the COAT trial and multi-locus sequence type (MSLT) was determined previously (5, 8). The strains used in this study are listed in Table 1. Strains were stored in glycerol stocks at −80°C. Strains from glycerol stocks were grown on YPD (yeast-peptone-dextrose) agar for 24-48 h. *C. neoformans* cells from agar plates were transferred to YPD broth and grown overnight at 30°C with shaking before being used in experiments.

**Table 1.**
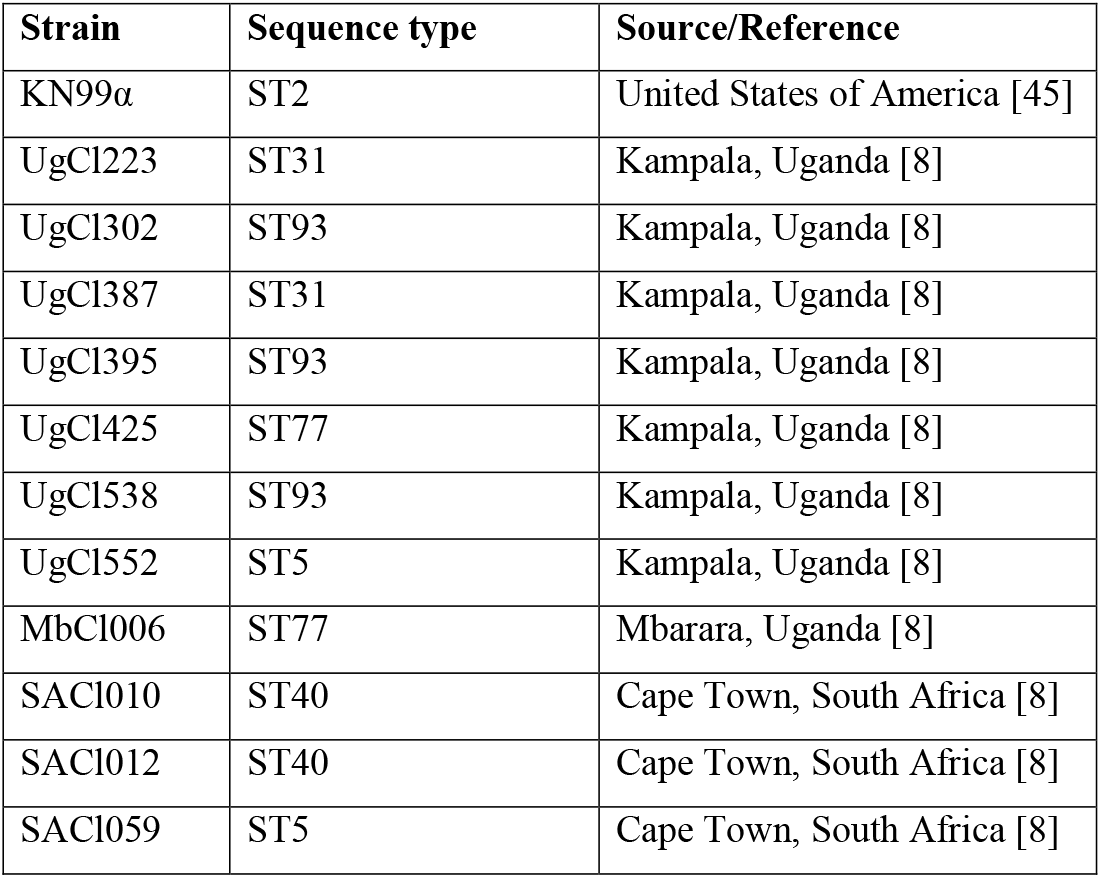
Strains used in this study.

| Strain | Sequence type | Source/Reference |
| --- | --- | --- |
| KN99α | ST2 | United States of America [45] |
| UgCI223 | ST31 | Kampala, Uganda [8] |
| UgCI302 | ST93 | Kampala, Uganda [8] |
| UgCI387 | ST31 | Kampala, Uganda [8] |
| UgCI395 | ST93 | Kampala, Uganda [8] |
| UgCI425 | ST77 | Kampala, Uganda [8] |
| UgCI538 | ST93 | Kampala, Uganda [8] |
| UgCI552 | ST5 | Kampala, Uganda [8] |
| MbCI006 | ST77 | Mbarara, Uganda [8] |
| SACI010 | ST40 | Cape Town, South Africa [8] |
| SACI012 | ST40 | Cape Town, South Africa [8] |
| SACI059 | ST5 | Cape Town, South Africa [8] |

### Mouse infection

*C. neoformans* strains were cultured overnight in YPD broth at 30°C. After incubation, *C. neoformans* cells were washed 3 times in sterile PBS, enumerated by hemocytometer and resuspended in sterile PBS at a concentration of 1 × 10^6^ cells/ml. Groups of 6-8 week-old female A/J mice (Jackson Laboratory, Bar Harbor, Maine) were anesthetized by intraperitoneal pentobarbital injection. 10 mice per strain were infected intranasally with 5 × 10^4^ cells in 50 μl PBS. Animals were monitored daily for morbidity and sacrificed when endpoint criteria were reached. Endpoint criteria were defined as loss of 20 % total body weight, loss of 2 grams in 2 consecutive days, or symptoms of neurological disease. Mice that survived to 150 days post-infection without exhibiting signs of disease were sacrificed and their tissues processed to determine lung, spleen and brain fungal burdens.

### Tissue burden analysis

Terminal lung, spleen and brain tissues were collected at the time of mouse sacrifice to determine organ fungal burden. Collected tissues were homogenized in 2 ml PBS. Serial dilutions of tissue homogenates were plated on YPD medium supplemented with 0.04 mg/ml chloramphenicol and incubated at 30°C. *C. neoformans* colonies were counted after 48 h of incubation. Data presented are representative of 2-8 mice per strain.

### Histopathology

Terminal lungs, spleen and brain were harvested from infected mice, fixed in 10% buffered formalin, paraffin-embedded, sectioned, and stained with H&E (hematoxylin and eosin). Tissue sections were examined for cell size and morphology by microscopy. Data presented are representative of 1-4 mice per strain.

### *In vivo* titan cell analysis

For a subset of infected mice, lungs were lavaged three times with 1.5 ml sterile PBS using a 20-gauge needle placed in the trachea. Cells in the lavage fluid were pelleted at 15,000 g and washed three times with PBS. Yeast cells were fixed with 3.7 *%* formaldehyde at room temperature for 40 min, washed 3 times with sterile PBS, and then resuspended in 200 μl sterile PBS. Cell body and capsule sizes were analyzed by microscopy. To visualize the yeast cells, a drop of india ink was added to the cell suspension on the slide and observed using a Zeiss Axioplan microscope (Axioimager, Carl Zeiss, Inc.). Cell body size was determined as the diameter of the yeast cell body and did not include the capsule. Data presented are representative of 1-3 mice per strain and a total of 200-400 *C. neoformans* cells per mouse were analyzed.

### *In vitro* titan cell analysis

To induce the formation of titan cells *in vitro*, two previously described methods were used (13, 14). For the first method (13), *C. neoformans* cells from a YPD agar plate were transferred to 10 ml YPD broth medium in a T25cm3 flask (TPP, Switzerland), grown at 30°C for 22 h with shaking (150 rpm), washed with sterile double distilled water and cell number enumerated with a hemocytometer. Aliquots of 1 × 10^6^ cells in 1 ml of minimal media (15 mM glucose, 10 mM MgSO4, 29.4 mM KH2PO4, 13 mM glycine, 3 μM Vit B1, pH5.5) were transferred into 1.5 ml Eppendorf tubes and incubated in a thermomixer at 30°C with shaking (800 rpm) for 48 hours. For the second method (14), yeast cells were grown overnight at 30°C, 150 rpm in 2 ml YNB without amino acids and supplemented with 2% glucose. Yeast cells from the overnight culture were washed twice with sterile PBS, cell number enumerated with a hemocytometer and resuspended in 0.5 ml PBS supplemented with fetal calf serum (FCS, ThermoFisher Scientific) to a final concentration of 1 × 10^3^ cells/ml in each well of a 24 well plate. The plates were then incubated at 37°C 5% CO2 for 48 h. After incubation, cell morphology was analyzed by microscopy (AxioImager, Carl Zeiss, Inc.). Cell body diameter was measured as described above. For *in vitro* titan cell analysis data presented are from 2 biological replicates per strain with 300 cells counted for each replicate.

### Capsule formation

Capsule formation was analyzed from *C. neoformans* cells generated from (a) *in vitro* titan cell formation as described above, and (b) *C. neoformans* cells grown in DMEM media supplemented with fetal calf serum (FCS). For capsule induction in DMEM, *C. neoformans* cells were grown overnight in YPD at 30°C with shaking (250rpm), washed with sterile PBS and counted with a hemocytometer. Yeast cells were then inoculated at a final concentration of 1x 106 cells/ml into DMEM media supplemented with 10% FCS and incubated at 37°C, 5% CO2 for 5 days. After incubation, yeast cells were fixed with 3.7% formaldehyde and analyzed by microscopy (Axioimager, Carl Zeiss, Inc.). Capsule width was defined as the difference between the diameter of the whole yeast cell (cell body and capsule) and the cell body diameter (no capsule) divided by two. Data presented are averages from a total of 100 *C. neoformans* cells per strain.

### Drug resistance assays

Drug resistance assays were as described previously (15). Briefly, a microdilution assay was performed according to CLSI guidelines using 2.5 × 10^3^ CFU/ml. Fluconazole and Amphotericin B were tested as 2-fold dilutions from 512 μg/ml to 0.0625 μg/ml and from 8 μg/ml to 0.0315 μg/ml, respectively, in a final volume of 200 μl per well. Spectrophotometric analysis of well turbidity at 600 nm was used to determine the minimum inhibitory concentration (MIC) for each strain. Plates were scanned in a Biotek Synergy H1 hybrid reader (Winooski, VT) prior to and after 72 h of incubation at 37°C. The amphotericin B MIC was defined as the drug concentration at which no growth was observed at 72 h (100% inhibition of growth). Fluconazole MIC was defined as a 50% reduction in growth (turbidity) compared to the no-drug control.

### Statistical analysis

T-test was used to compare clinical parameters between patients who lived and those who died. A Fisher exact test was used to analyze the association between survival in mice and survival in human patients. P-values < 0.05 were considered significant.

## Results

### *C. neoformans* isolates with the same sequence type were associated with both mortality and patient survival

The association between clinical outcome and *C. neoformans* MLST-based sequence type (ST) is controversial, with some studies showing differences in patient clinical outcomes associated with specific sequence types and others showing no correlation (5, 7–9, 16). To explore this phenomenon, we examined individual patient mortality/survival in HIV+ cryptococcal meningitis patients (enrolled in the Cryptococcal Optimal ART Timing (COAT) clinical trial) that were infected with *C. neoformans* strains of the same sequence type (Fig 1A). In this cohort, 11 sequence types were observed in only one patient and were cumulatively analyzed as “unique ST” types. We observed a combination of patient death within 10-weeks (early mortality) and patient survival for eight of the sequence types (ST5, ST31, ST77, ST65, ST93, ST2, ST206 and ST187). No patients survived infections with ST40, whereas all patients survived the ST23 and ST71 infections (Table 2). While ST40 and ST23 patient mortality/survival were examples of the two ends of the spectrum, patient mortality (# patients with mortality due to cryptococcosis / total patients) differed across the sequence types (Figure 1A and Table 2). For example, ST187 had a high rate of patient mortality (i.e. low survival) whereas ST5 had a lower rate of patient mortality. Similar patient mortality/survival was observed with the unique sequence types.

**Figure 1.**
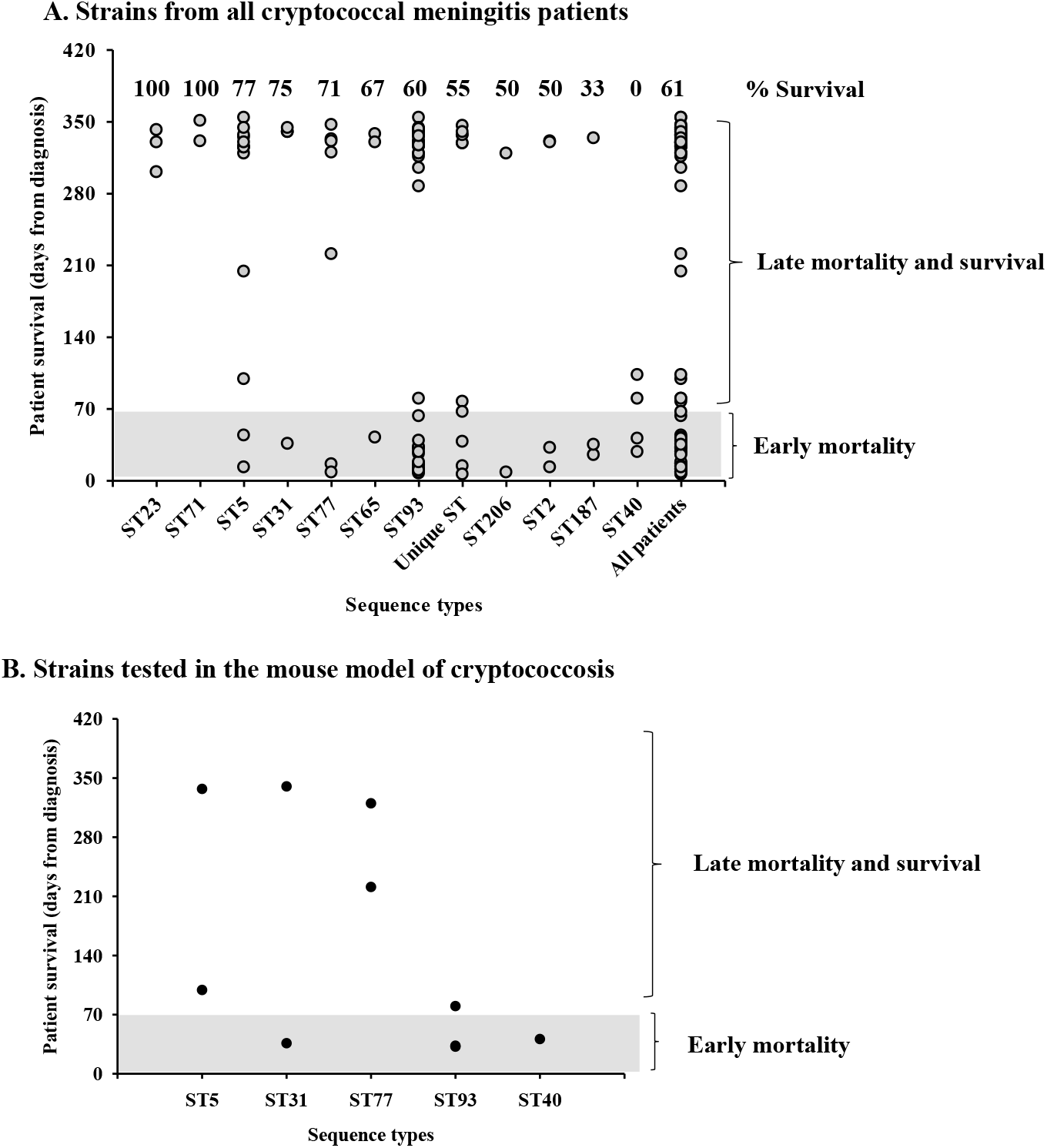
*C. neoformans* sequence types and human mortality. *C. neoformans* clinical strains were isolated from the CSF of HIV-CM patients, grown on YPD agar media at 30°C and stored at −80°C before their sequence types were identified. Early mortality: died within 10 weeks from diagnosis, late mortality: died after 10 weeks, survival: survived the infection. Unique ST; sequence types with only one patient.

**Table 2.** HIV-CM patient infection outcome.

|  | Sequence types |  |  |  |  |  |  |  |  |  |  |  |  |
| --- | --- | --- | --- | --- | --- | --- | --- | --- | --- | --- | --- | --- | --- |
|  | ST23 | ST71 | ST5 | ST31 | ST77 | ST65 | ST93 | Unique<br>ST | ST206 | ST2 | ST187 | ST40 | All<br>patients |
| Early mortality<br>(within 10 weeks) | 0 | 0 | 2 | 1 | 2 | 1 | 19 | 4 | 1 | 2 | 2 | 2 | 36 |
| Late mortality (after<br>10 weeks) | 0 | 0 | 1 | 0 | 0 | 0 | 1 | 1 | 0 | 0 | 0 | 2 | 5 |
| Survival | 3 | 2 | 10 | 3 | 5 | 2 | 30 | 6 | 1 | 2 | 1 | 0 | 65 |
| Total | 3 | 2 | 13 | 4 | 7 | 3 | 50 | 11 | 2 | 4 | 3 | 4 | 106 |
| Survival % | 100.0 | 100.0 | 76.9 | 75.0 | 71.4 | 66.7 | 60.0 | 54.5 | 50.0 | 50.0 | 33.3 | 0.0 | 61.3 |

These data show that the majority of sequence types had patients that succumbed to the *C. neoformans* infection and patients that survived the infection. There are a number of parameters, both patient and fungal, that could explain this difference in patient survival within the same sequence type. We investigated factors that could impact whether a patient survived or succumbed to cryptococcal infection with the same sequence type.

### Association between HIV disease and mortality/survival

*C. neoformans* infections are predominantly observed in patients with compromised immune function. To test the hypothesis that differences in the underlying HIV infection account for the observed differences in mortality/survival in the various sequence types, we examined CD4+ counts, cerebrospinal fluid white blood cell count (CSF_WBC), and HIV viral load in patients that died from cryptococcosis within 10-weeks vs. those that survived (Fig 2A-C). ST93 was the most prevalent sequence type, with 60 patients, and the only sequence type with enough patients to perform statistical analyses. Therefore, we also analyzed HIV viral load, CSF_WBC, and CD4+ cell count in the cohort of patients infected with ST93 strains alongside analyzes for all patients (Fig 2D-F). The number of CD4+ cells, CSF white blood cells and the HIV viral load was similar in patients that lived and those that died (p-value > 0.05). Similar results were observed when considering only the strains that were tested in the mouse model of cryptococcosis (Supplemental Figure 1).

**Figure 2.**
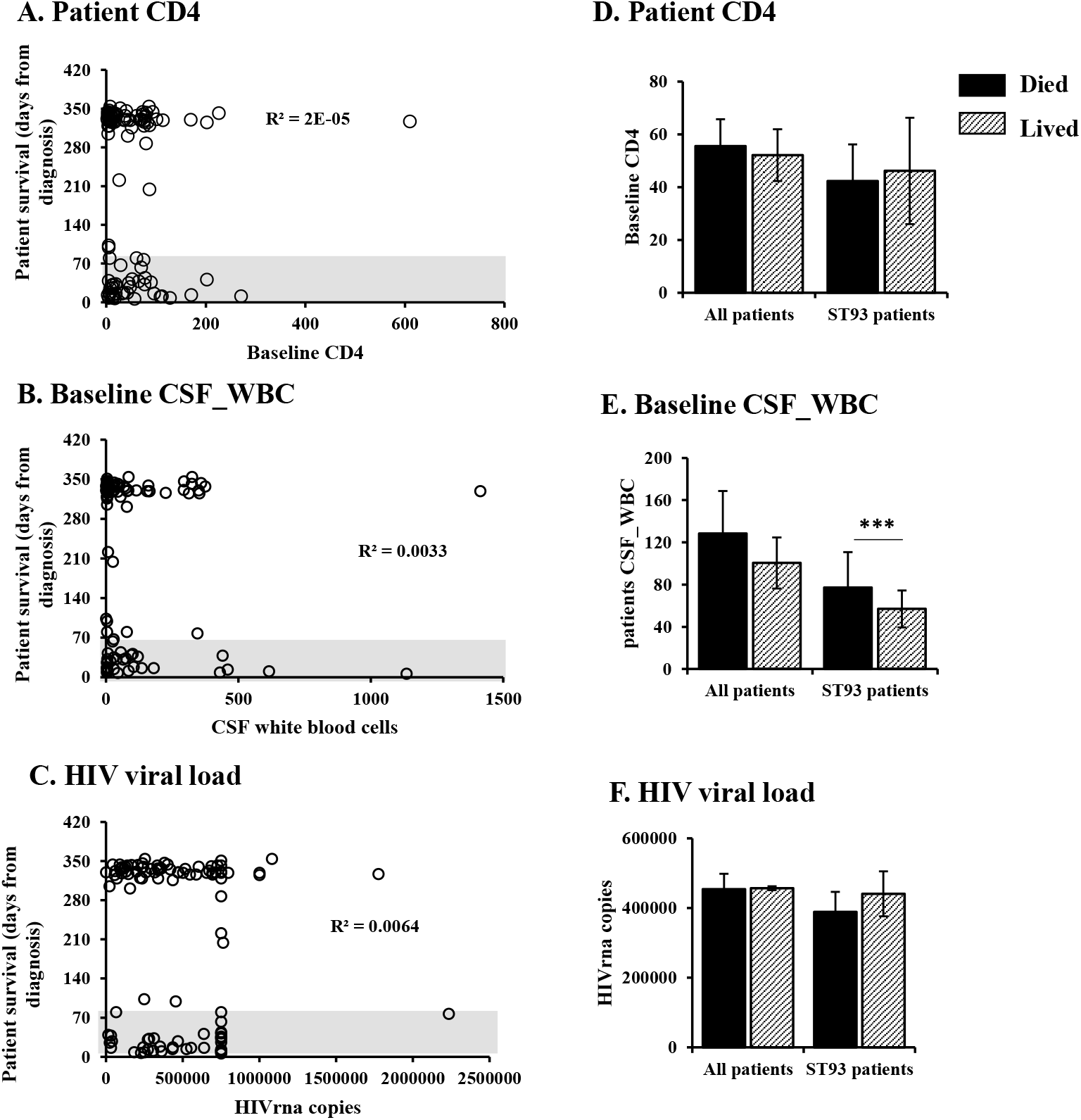
Human mortality is not determined by patient parameters. Clinical outcome was compared to baseline CM patient characteristics including CD4+ cells (A, D), white blood cell counts in the CSF (B, E), and HIV viral load (C, F). Graphs A-C show individual patient parameters plotted against survival from diagnosis. Gray bar indicates mortality within 10-weeks. Survival beyond 280 days is indicated as the discharge date from the clinical trial. Graphs D-F show patient parameters when classified as died (mortality) by 10 weeks post-diagnosis or lived past 10 weeks post-diagnosis. Error bars represent standard error of the mean. CD4 cells: CD4+ T helper cells, CM: cryptococcal meningitis, CSF: cerebrospinal fluid, HIV: human immunodefiency virus, RNA: ribonucleic acid, ST93: sequence type 93, WBC: white blood

Although the CSF_WBC counts did not differ between patients who died and those that survived the infection in the total cohort, patients who survived the ST93 infection had lower CSF_WBC counts compared to patients who died from the ST93 infections (Fig 2E, p-value < 0.05). These data suggest that observations at the population level do not always represent individual sequence types, and that higher WBC count at the time of presentation is associated with mortality in the ST93 infected patients.

### Human mortality is associated with fungal burden in the CSF

To explore the hypothesis that fungal factors impact mortality/survival in the various sequence types, we analyzed the initial CSF fungal burden (CSF_CFUs), the amount of cryptococcal antigen in the CSF (CrAg) and the rate of fungal clearance in the patient CSF (EFA). Our analysis revealed that these fungal parameters partially correlate with patient survival (Fig 3). Looking at all patients individually, we see that patients with comparable cryptococcal CFUs have different clinical outcome (Fig 3A). However, lower CFUs were associated with survival in the ST93 infected patients (Fig 3D). We observed similar trends for the amount of cryptococcal capsular antigen detected in the patient CSF and the rate of fungal clearance (Fig 3B, E and C, F) with higher antigen levels and lower clearance associated with mortality in ST93 genotypes but not in the entire population. These data suggest that fungal parameters such as the ability to multiply inside the host, resistance to treatment and amount of antigen shedding by *Cryptococcus* cells play a role in infection outcome in a sequence type dependent manner.

**Figure 3.**
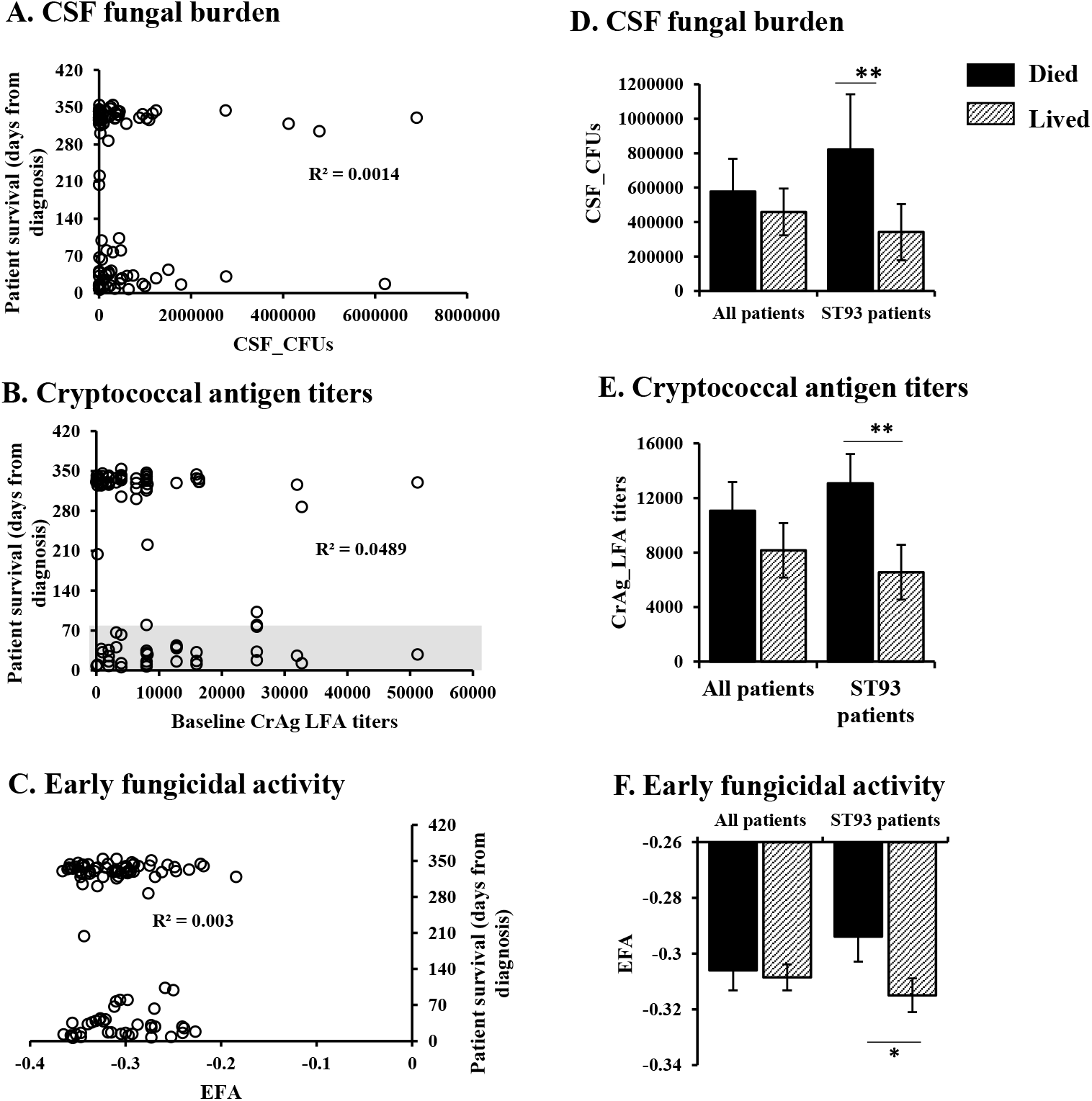
Patient survival is partly associated with fungal parameters. Clinical outcome was compared to fungal parameters including CSF-fungal burden determined by colony forming units (CFUs) (A, D), cryptococcal antigen titers in the CSF (B, E) and rate of fungal clearance determined by the early fungicidal activity (EFA) (C, F). Graphs A-C show individual patient parameters plotted against survival from diagnosis. Graphs D-F show patient parameters when classified as died (mortality) by 10-weeks postdiagnosis or lived (survived) past 10-weeks post-diagnosis. Error bars represent standard error of the mean. CFU: colony forming unit, CSF: cerebrospinal fluid, CrAg LFA: cryptococcal antigen lateral flow assay, EFA: early fungicidal activity, ST93: sequence type 93. *p< 0.05, **p< 0.01.

When only the patients infected with strains that were tested in the mouse model of cryptococcosis were considered, only the fungal burden in the CSF was associated with virulence (Supplemental Fig 2 A, D). The cryptococcal antigen detected in the patient CSF as well as the rate of fungal clearance were similar between the patients infected with high, intermediate and low virulence strains (Supplemental Fig 2 B, E and C, F).

### *C. neoformans* virulence in human patients correlates with virulence in a mouse model of cryptococcosis

After observing that both human and fungal parameters were partially associated with patient survival, we determined whether differences in *Cryptococcus* virulence in humans could be recapitulated in the mouse inhalation model of cryptococcosis. We selected paired strains - one from a patient that died and one from a patient that lived – from ST5, ST31, and ST77 for analysis in the mouse model (Fig 1B). In addition, we analyzed two high-mortality ST40 strains, two high-mortality ST93 strains, and one late-mortality ST93 strain (Fig 1B). Mice infected with the clinical strains showed variation in disease progression in the mouse inhalation model (Fig 4). Based on mouse survival, the clinical strains were divided into three groups: high, intermediate and low virulence strains (Figures 4A-C). The high virulence strains were defined as strains that caused 80% mortality within 37 days post-infection (Fig 4A). The intermediate virulence strains had mortality between 69-129 days post-infection (Fig 4B). The low virulence group consisted of three strains; mice infected with two strains, UgCl552 and SACl010, showed no overt signs of the disease at 150 days post-infection, and the third strain, UgCl223 had reduced virulence where 60% of mice survived through 150 days post-infection (Fig 4C). By histologic examination of lungs, no cryptococci were identified in UgCl552 or SACl010, however UgCl223 had a low to moderate number of organisms within the lung and one of two mice examined had infiltration of small cryptococcal organisms into the brain with necrosis in affected areas.

**Figure 4.**
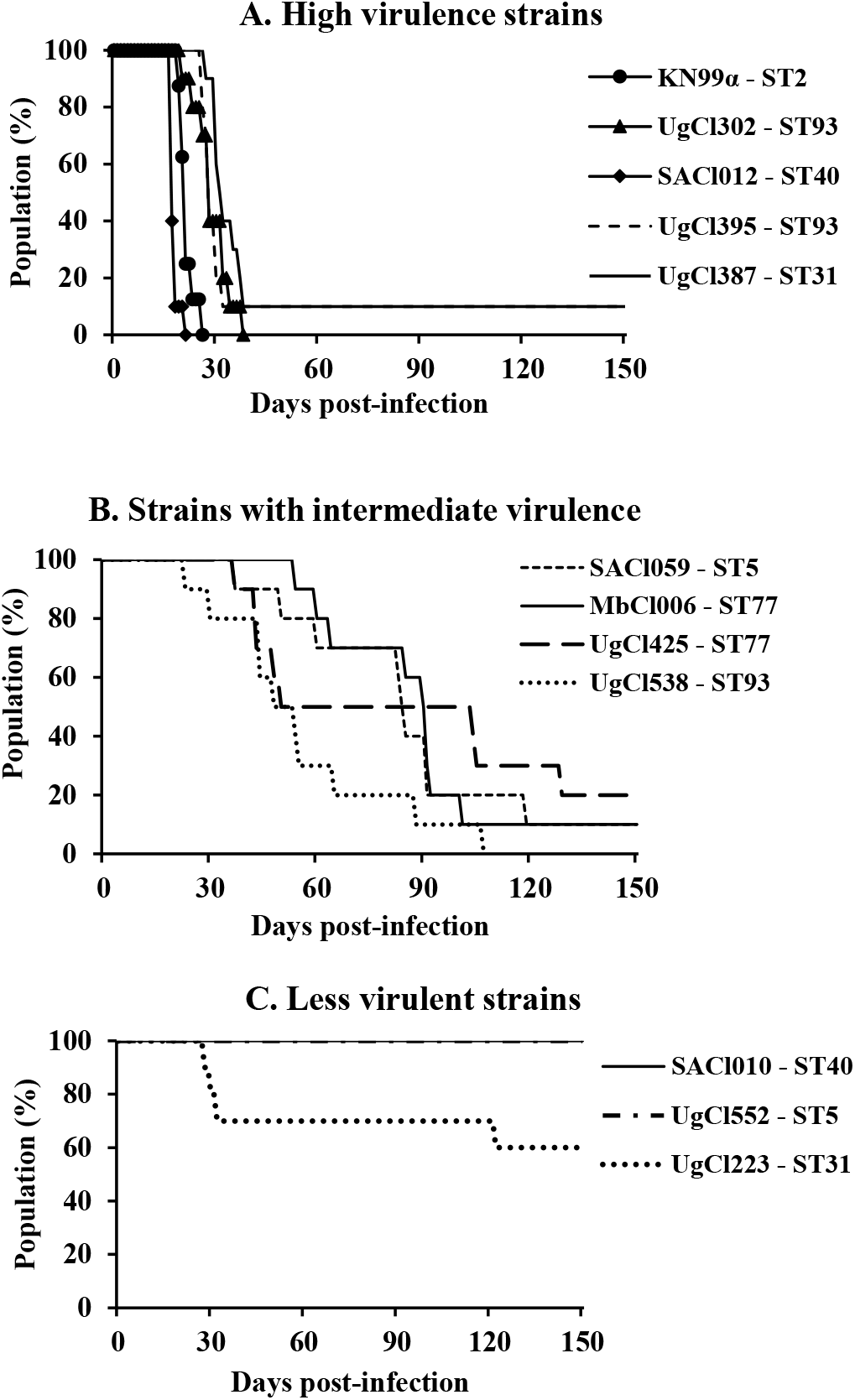
*C. neoformans* clinical strains show differential virulence in mice. Groups of ten 6-8 week old A/J mice were infected intranasally with 5 × 10^4^ cells from *C. neoformans* clinical strains isolated from the CSF of HIV-CM patients. Progression to severe morbidity was monitored for 150 days and mice were sacrificed when end point criteria were reached. A) high virulence strains, B) strains with intermediate virulence, C) low virulence strains.

We next compared the death/survival of patients and mice infected with the same *Cryptococcus* strain (Figure 5 and Table 3). All but two of the isolates associated with early mortality in humans showed high virulence in mice (Figure 5A). Strains SACl010 and UgCl552 had low virulence in mice but had an early mortality in humans. Histological examination of lungs from 2 mice infected with SAC1010 revealed low numbers of cryptococci within alveoli that were eliciting a strong pulmonary inflammatory response. Interestingly, while the human patient infected with SACl010 died 41 days after diagnosis, the clinical records indicated that the patient died from Immune Reconstitution Inflammatory Syndrome (IRIS) and not cryptococcosis. IRIS is defined as detrimental inflammatory responses despite fungal clearance (e.g. negative cultures) (17), consistent with the inflammation observed in the mouse model. The patient infected with strain UgCl552 cleared the initial infection but then died at 99 days post-diagnosis of an unknown cause at home; again, consistent with the mouse model showing low virulence of the *Cryptococcus* strain.

**Figure 5.**
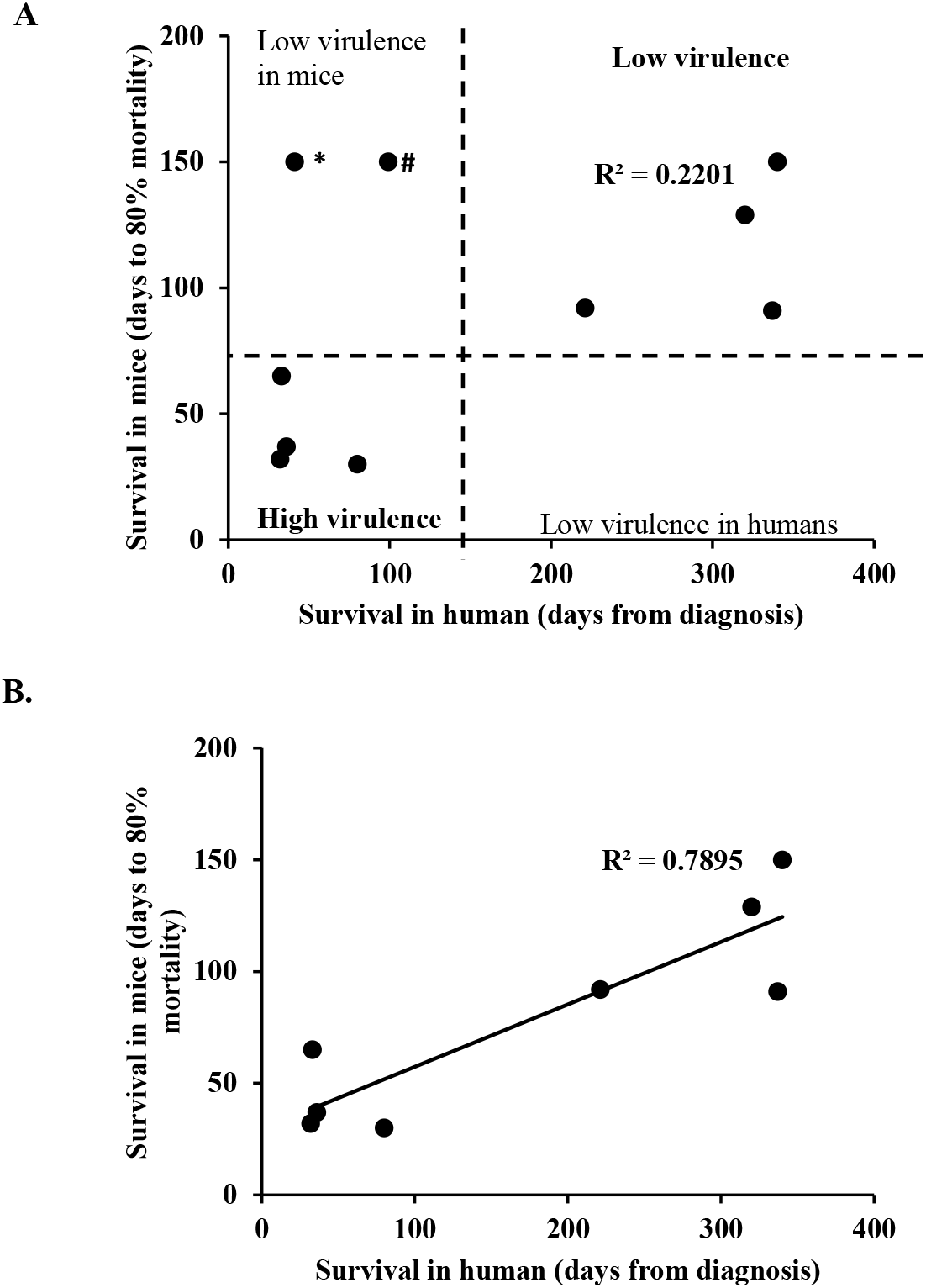
Infection outcome in human patients is associated with disease outcome in mice. Clinical outcome in human patients was compared to infection outcome in mice infected with the same *C. neoformans* strain. A) Correlation between human and mice survival for all 10 clinical strains. B) Correlation between human and mice survival when outliers (SACl010 and UgCl552 strains) were removed. * The SACl 010 strain was associated with low virulence in mice but the human patient died of IRIS at 41 days post-diagnosis. # The patient infected with UgCl552 died at home after clearing the initial *C. neoformans* infection. The cause of death was not determined.

**Table 3.** Association between clinical strain virulence in human and mice.

|  |  |  | Human infections |  | Mice infection |  |
| --- | --- | --- | --- | --- | --- | --- |
|  | Strain name | Sequence type | Mortality (days from diagnosis) | Outcome | Mortality (Days to 80% mortality) | Outcome |
| High virulence | SACI012 | ST40 | NA | NA | 18 | All died |
| | KN99α | ST2 | NA - WT | NA - WT | 21 | All died |
|  | UgCI395 | ST93 | 80 | Died | 30 | Died (9/10)<sup>1</sup> |
|  | UgCI302 | ST93 | 32 | Died | 32 | Died (9/10) |
|  | UgCI387 | ST31 | 36 | Died | 37 | Died (9/10)<sup>2</sup> |
| Intermediate virulence | UgCI538 | ST93 | 33 | Died | 65 | All died |
|  | SACI059 | ST5 | - | Survived | 91 | Died (9/10)<sup>3</sup> |
|  | MbCI006 | ST77 | 221 | Died | 92 | Died (9/10)<sup>4</sup> |
|  | UgCI425 | ST77 | - | Survived | 129 | Died (8/10)<sup>5</sup> |
| Low virulence | UgCI223 | ST31 | - | Survived | End of experiment | Survived |
|  | UgCI552 | ST5 | 99 | IRIS | End of experiment | Survived |
|  | SACI010 | ST40 | 41 |  | End of experiment | Survived |

<sup>1</sup> Nine out of 10 mice died, with remaining mouse sacrificed after 150 days, terminal CFUs of surviving mouse were $6.4 \times 10^5$ cells in the lungs and $4 \times 10^2$ in the brain tissues
<sup>2</sup> Nine out of 10 mice died, with remaining mouse sacrificed after 150 days, terminal CFUs of surviving mouse were $7 \times 10^5$ cells in the lungs, no *Cryptococcus* cells detected in the brain
<sup>3</sup> Nine out of 10 mice died, with remaining mouse sacrificed after 150 days, terminal CFUs were $6 \times 10^5$ cells in the lungs, no *Cryptococcus* detected in the brain
<sup>4</sup> Nine out 10 mice died, with remaining mouse sacrificed after 150 days, terminal CFUs were $6 \times 10^6$ cells in the lungs, and $6 \times 10^3$ cells in the brain
<sup>5</sup> Eight out of 10 mice died, with remaining mice sacrificed after 150 days, terminal CFUs were $4 \times 10^5$ cells in the lungs of each mouse and $4 \times 10^4$ cells in the brain of one mouse and $3.6 \times 10^4$ cells in the brain of the second mouse
NA: patient information not available, NA-WT: patient information not available, wild type laboratory control strain, NI: not identified at the time of death, patient died at home.

All strains associated with late mortality or survival in human patients showed reduced virulence in mice (Figure 5 and Table 3). No strain had high virulence in mice but low virulence in humans, providing further support for the association between human and mouse virulence. When all 10 strains (including the outliers SACl010 and UgCl552) were considered, a trend between human and mouse survival was observed (Figure 5A, R^2^= 0.2201). However, when both outliers are removed, the association between human and mouse survival is robust (Figure 5B, R^2^= 0.7895). These data show human and mouse mortality are similar when infected with the same *C. neoformans* strain, although unrelated human causes of mortality can influence the human clinical outcome.

### Mouse virulence is associated with fungal burden

To further investigate the factors that influence infection outcome in mice, we compared mouse survival with tissue fungal burden at sacrifice. High virulence strains were associated with high fungal burdens while intermediate and low virulence strains correlated with low CFUs in lungs of infected mice (Fig 6A). Similar trends were observed for fungal burdens in the spleen, but differences were not statistically significant (Fig 6B). We observed similar terminal fungal burdens in the brains of mice infected with high and intermediate virulence strains, and lower fungal burdens in the mice infected with low virulence strains sacrificed at 150 days post-infection (Fig 6C). These data suggest that both high and intermediate virulence strains can disseminate to the brain, the only difference being the time it takes to reach the brain tissue (Fig 6C and Table 3). Histologic examination of lung and brain from two intermediate strains UgC1425 and SAC1059 revealed moderate to large numbers of cryptococci and cryptococcoma formation with a moderate to strong inflammatory response.

**Figure 6.**
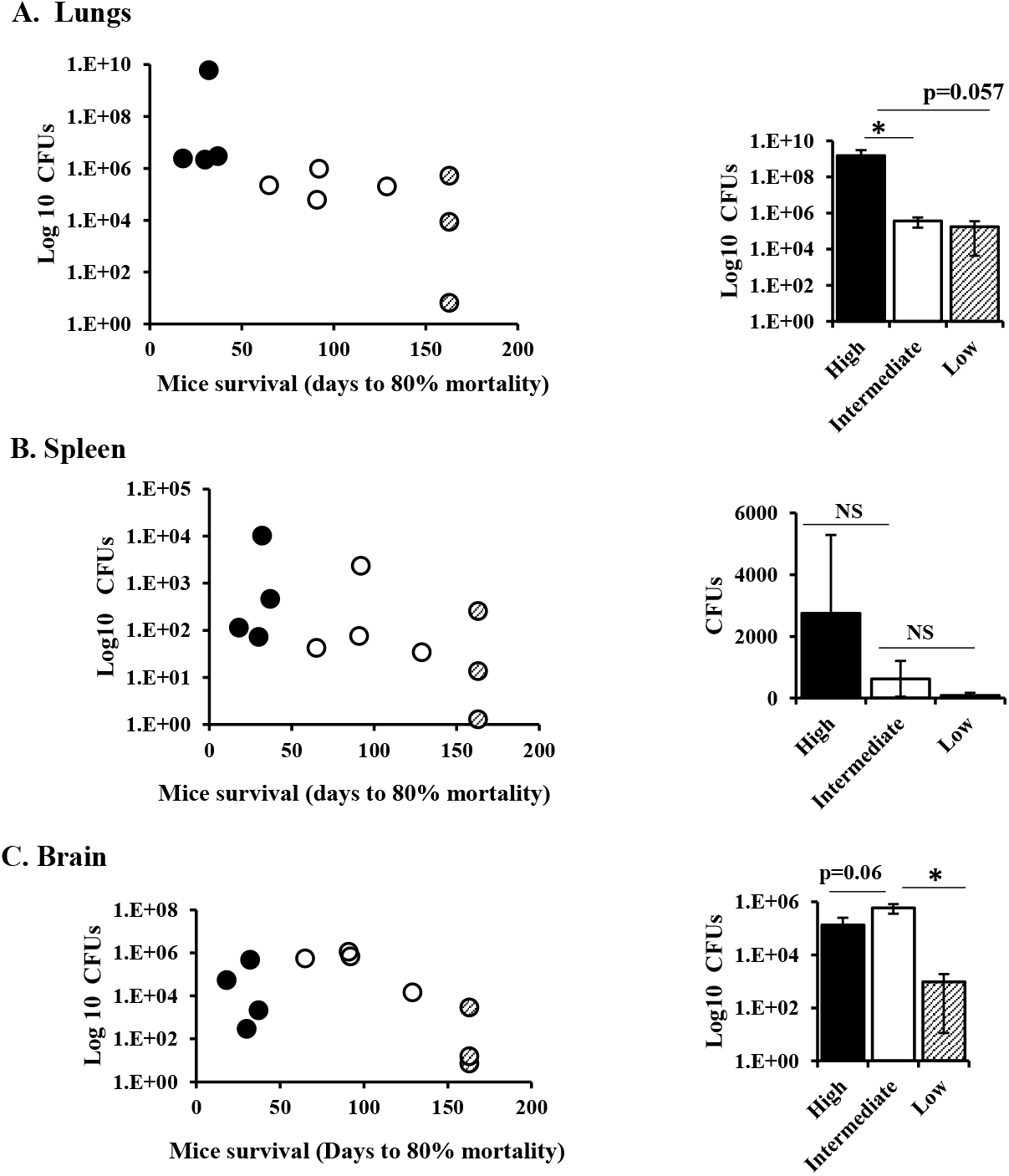
Mouse survival correlates with organ fungal burden. Groups of ten 6-8 week old A/J mice were infected intranasally with eleven *C. neoformans* clinical strains and a wild type laboratory strain. Progression to severe morbidity was monitored for 160 days, and mice were sacrificed at 160 days or when end point criteria were reached. Lungs (A), spleen (B) and brain (C) were harvested, homogenized and serial dilutions were plated on agar plates to determine tissue fungal burdens. Strains were classified in three groups as high, intermediate and low virulence strains as determined in the mice survival experiment. Left graphs show individual *C. neoformans* strains plotted as tissue fungal burdens against time to 80 % mortality of infected mice. Graphs on the right show average CFUs of high, intermediate and low virulence group strains. Error bars represent standard error of the mean. CFU: colony forming unit, NS: not statistically significant, PBS: phosphate buffered saline. * p< 0.05.

Mice infected with low virulence strains showing no overt disease were sacrificed at 150 days post-infection. These mice had moderate CFUs in the lungs but low CFUs in the spleen and brain, suggesting that the low virulence of these strains was due to reduced fungal dissemination (Fig 6 B-C). In addition, we observed differences in the disease progression/pathogenesis in mice infected with the three low virulence strains. One of the low virulence strains (UgCl223) had high terminal lung CFUs (1X 10^7^) and very few CFUs in the spleen (10) and brain (10). Histologic examination of lung and brain from 2 additional animals revealed low numbers of cryptococci in the lung with moderate inflammation, and one mouse had small cryptococci infiltrating the meninges, scattered in small areas of necrosis within the neuropil. These data suggest that UgCl223 can multiply in the lungs and may disseminate to the brain but may not remain viable in the brain. Most mice infected with the other two low virulence strains (SACl010 and UgCl552) cleared the infection. Seven out of nine mice infected with SACl010 had no CFUs in the lung, spleen or brain tissues at 150 days post-infection. Five out of nine mice infected with UgCl552 had no CFUs in the lungs and only one mouse with lung CFUs had detectable CFUs in the spleen or brain.

### *C. neoformans* virulence did not correlate with known virulence factors in *in vitro* conditions

After observing that infection outcome was strain-specific in both human and mouse infection, we tested whether production of virulence factors previously shown to be associated with *C. neoformans* pathogenesis (18–23) were associated with survival in our study. Virulence factors tested include the ability to growth at high temperature (37°C), capsule formation, and the ability to form titan cells both *in vitro* and *in vivo*. Virulence did not correlate with the ability to grow *in vitro* in either nutrient-rich and limited nutrient conditions, at low and high temperatures (30°C and 37°C) (Supplemental Figure 3). Using two previously described methods (13, 14), we found that virulence was also not associated with the ability to form titan cells *in vitro* (Supplemental Table 1). However, we observed a trend where low virulence strains formed more titan cells than high and intermediate virulence strains when grown in DMEM media supplemented with serum (Table 4). Finally, we also analyzed *in vitro* capsule formation. While capsule varied across the strains, there was no difference in the size of the capsule between low and high virulence strains (Supplemental Figure 4).

**Table 4.** Cellular phenotypes and growth in the presence of cell wall stressors.

|  |  | Titan cells in DMEM+FCS (%) | Capsule size in DMEM+FCS (μm) | Fluconazole heteroresistance (μg/ml) | Growth on Congo Red | Growth on CFW | Growth on Caffeine |
| --- | --- | --- | --- | --- | --- | --- | --- |
| High resistance | KN99α | 0 | 3.8 | 16 | ++++ | ++++ | +++ |
|  | SACI012 | 0 | 2.5 | 8 | +++ | ++++ | ++++ |
|  | UgCI302 | 6 | 5.0 | 32 | ++++ | ++++ | ++++ |
|  | UgCI395 | 0 | 7.5 | 32 | +++ | ++++ | ++++ |
|  | UgCI387 | 0 | 7.4 | 32 | +++ | ++++ | +++ |
| Intermediate resistance | UgCI425 | 0 | 3.4 | 32 | ++++ | +++ | +++ |
|  | UgCI538 | 1 | 5.8 | 16 | +++ | ++++ | ++++ |
|  | SACL059 | 0 | 3.3 | 16 | ++++ | +++ | ++++ |
|  | MBCL006 | 3 | 4.8 | 16 | ++++ | ++++ | ++++ |
| Low resistance | UgCI223 | 1 | 3.1 | 16 | ++++ | +++ | ++++ |
|  | UgCI552 | 4 | 3.1 | 4 | - | - | - |
|  | SACL010 | 10 | 6.3 | 16 | - | - | ++ |

“-” No growth observed, “+” number of dilutions that grew on agar media by spotting assay

We next compared the susceptibility of the clinical strains to the antifungal drugs fluconazole and amphotericin B, which were used to treat the CM-patients in our cohort. The *in vitro* susceptibility to the antifungal drugs fluconazole and amphotericin B was not associated with strain virulence (Supplemental Table 2). Next, we analyzed the levels of heteroresistance to fluconazole, a factor that was previously associated to *C. neoformans* virulence (24), as well as the ability to grow in the presence of cell wall stressors calcofluor white (CFW), congo red and caffeine. We observed a trend where high virulence strains manifested heteroresistance at high levels of fluconazole and low virulence strains developed heteroresistance at low levels of fluconazole (Table 4). However, not all high virulence strains had high levels of fluconazole heteroresistance. Two highly virulent strains, SACl012 and KN99α, manifested fluconazole heteroresistance at 8 and 16 μg/ml respectively, while the other three high virulence strains showed heteroresistance at 32 μg/ml (Table 4). In addition, high virulence strains grew better in the presence of cell wall stressors compared to two of the three low virulence, SACl010 and UgCl552 (Table 4). However, the low virulence strain UgCl223 did not show any growth defect in any of the tested conditions (Table 4). Our *in vitro* assays did not identify a single condition or a virulence-determining factor that exclusively distinguishes our high and low virulence strains, but there was a trend where most of low virulence strains grew slower or did not grow in the presence of various stresses.

## Discussion

In this study, we showed that patients infected with *C. neoformans* strains of the same sequence type could have different clinical outcomes. Using a mouse inhalation model of cryptococcosis, we showed that these differences in virulence were strain-specific – with similar mortality rates observed in both humans and mice infected with the same strain. In both humans and mice, we identified an association between increased fungal burden and early mortality, but no single *in vitro* virulence factor could explain the observed *in vivo* differences between closely related strains.

We sought to identify factors associated with infection outcome, both in humans and infected mice, which could explain the observed differences in mortality between *C. neoformans* strains with the same sequence type. We found that *C. neoformans* growth *in vivo* was associated with mortality rate in mice and humans. Our observations are in accord with previous clinical trials showing that CSF fungal burden and the rate of fungal clearance were associated with patient mortality (25–28). However, underlying human immune deficiencies did not correlate with patient outcome. Some patients died, others survived the infection, while having similar CD4+ cell counts, HIV viral load and white blood cell numbers in their CSF. These data are also consistent with previous observations that baseline CD4+ cells, CSF white blood cell counts, and HIV viral load do not influence the mortality of HIV-CM patients (29–31). Taken together, these data show that differences in patient outcome are due, in part, to differences in the strains of *C. neoformans* with which the patients are infected. A previous study from our group showed that differences in *C. neoformans* genotype influence patient immune response and clinical outcome in HIV-CM patients (5). To further explore this phenomenon, we examined the impact of sequence type on patient outcome. We found that patients infected with *C. neoformans* isolates of the same sequence type had differential clinical outcome, some patients survived while others died from the infection.

To determine whether these differences in clinical outcome where inherent to the individual *C. neoformans* strains, we tested the virulence of a subset of the clinical isolates in the mouse inhalational model of cryptococcosis. We showed a robust association between patient and mouse mortality for the majority of *C. neoformans* strains tested. These data not only show that the mouse model accurately recapitulates human disease, but also strongly suggest that a large proportion of patient mortality is due to inherent differences between the infecting *C. neoformans* strains.

Several factors such as cellular phenotypes and stress response mechanisms have been associated with *C. neoformans* virulence and pathogenesis in the mouse model of cryptococcosis (18, 21, 22). In addition, a few studies showed that pleomorphism and the production of some virulence factors under *in vitro* conditions correlated with infection outcome in human patients (7, 23, 32). Therefore, we tested whether the ability to produce known virulence factors, as well as other phenotypes previously associated with pathogenesis, correlate with the virulence level of the subset of *C. neoformans* clinical isolates used in our study. The dominant virulence factors in *C. neoformans* are high temperature growth and capsule formation (33, 34). High and low virulence strains grew similarly at high temperature, both in nutrient-rich and low nutrient conditions. We observed similar patterns in capsule formation where high and low virulence strains produced capsule of similar size. However, the size of the capsule formed during infection was significantly greater that the capsule induced *in vitro* suggesting differences in capsule structure, composition and/or function (35–37). This might explain our observations and those from other studies (38–40) that capsule size *in vitro* does not correlate with *C. neoformans* virulence. Next, we tested whether increased virulence was associated with increased resistance to two antifungal drugs, fluconazole and amphotericin B, used to treat patients in our cohort. *In vitro* susceptibility to fluconazole and amphotericin B also did not correlate with virulence of our clinical strains. The lack of correlation between infection outcome and the susceptibility to antifungal drugs in *in vitro* assays can be explained by physiological differences between *in vitro* conditions and *in vivo* environments in which *C. neoformans* cells have to survive (41). We also tested the ability to form titan cells, a phenotype that has been associated with *C. neoformans* virulence/pathogenesis (18, 19, 42). Using recently described techniques to induce titan cells *in vitro* (13, 14), we found that the ability to form titan cells *in vitro* did not correlate with the degree of virulence of our clinical strains. Similar results were observed with *in vivo* titan cell formation in the mouse model.

Under *in vivo* conditions, *C. neoformans* cells have to withstand multiple host defense mechanisms. Our data show that, individually, the *in vitro* analyses we tested were unable to distinguish high versus low virulence strains. Thus, no single *in vitro* assay/condition can be used as a proxy for *in vivo C. neoformans* virulence. However, we did observe a trend when comparing across the various *in vitro* assays (Table 4). Increased virulence was associated with heteroresistance at high levels of fluconazole, ability to grow in the presence of cell wall stressors, and inability to form titan cells in DMEM and serum media. In contrast, strains with low virulence were unable to grow in the presence of cell wall stressors, had low fluconazole heteroresistance and formed more titan cells in DMEM and serum. Most mice infected with strains that were unable to grow in the presence of multiple stresses *in vitro* cleared the infection and the strains did not disseminate to the brain, showing that growth defects *in vitro* in response to multiple stresses might predict inability to multiply *in vivo* during infection. These observations suggest that a panel of *in vitro* stresses could be developed to differentiate between high and low virulence *C. neoformans* strains. Future studies that incorporate a large number of clinical isolates are necessary to determine whether an appropriate panel of *in vitro* stresses can be identified to accurately predict *in vivo* virulence. If confirmed, this could be a valuable tool in clinical laboratories to help in the early identification and follow up of patients infected with high virulence strains and at high mortality risk.

An alternative explanation for differences in *in vivo* virulence that do not correlate with single *in vitro* assays is that the observed differences are due to novel virulence factors. *In vivo* studies with evolutionarily closely related strains, such as those presented here with identical ST types, need to be combined with genomic analyses to identify novel genes that are critical *in vivo*. Our observation that strains with identical sequence type can have dramatically different virulence in both humans and the mouse model suggest that multi-locus sequence type does not have sufficient resolution to identify the underlying differences between strains – whole genome sequencing of closely related strains, and at the population level, may be required to identify these *in vivo* virulence factors.

In summary, our study had two major findings. First, the mouse inhalation model of cryptococcosis accurately recapitulates human infection. Thus, this model can be used to explore how differences between *C. neoformans* clinical isolates impact human disease outcome. Second, *C. neoformans* isolates from the same sequence type were associated with different clinical outcomes. The association between sequence type and clinical outcome has been contentious, with some patient cohorts showing no association and some with robust associations (5, 7, 43, 44). Our data show that belonging to the same lineage or sequence type does not necessary mean *C. neoformans* strains will have a comparable degree of virulence. The virulence-determining factor(s) is not the lineage/sequence type of the *C. neoformans* strain, but instead other genotypic or phenotypic characteristics specific to individual isolates within the sequence type. Future studies investigating the relationship between individual *C. neoformans* genotypes and virulence are needed to fully understand pathogen-associated factors that influence *in vivo* virulence and ultimately clinical outcome.

## Acknowledgements

This research was supported by NIH grants R01AI134636 (KN), R21AI122352 (KN), and U01AI089244 (DRB). LM received support from the NIH Fogarty International Center (R25TW009345) and the University of Minnesota Center for Translational Science Institute (UL1TR000114). We thank the University of North Carolina Lineberger Comprehensive Cancer Center Animal Histopathology Core Facility for their help with this studies

---

**Supplemental Table 1.** Titan cell formation.

|  |  | *In vitro* |  | *In vivo* |
| --- | --- | --- | --- | --- |
|  |  | Protocol-1 | Protocol-2 |  |
| High virulence | KN99α | + | + | + |
|  | UgCI302 | - | - | + |
|  | UgCI387 | - | - | + |
|  | UgCI395 | - | - | + |
|  | SACI012 | - | - | - |
| Intermediate virulence | UgCI425 | - | - | + |
|  | UgCI538 | - | - | + |
|  | SACI059 | - | - | + |
|  | MbCI006 | - | - | + |
| Low virulence | UgCI223 | - | - | + |
|  | UgCI552 | + | - | + |
|  | SACI010 | + | - | + |

+: presence of titan cells, -: no titan cell observed

**Supplemental Table 2.** Minimum Inhibitory Concentration (MIC) for fluconazole and amphotericin B is not associated with strain virulence.

|  | Strain | Fluconazole<br>MIC | Amphotericin<br>B MIC |
| --- | --- | --- | --- |
| High virulence | KN99α | 1 | 0.5 |
|  | SACI012 | 1 | 0.25 |
|  | UgCI302 | 1 | 1 |
|  | UgCI395 | 2 | 0.5 |
|  | UgCI387 | 0.5 | 0.5 |
| Intermediate<br>virulence | UgCI425 | 2 | 1 |
|  | UgCI538 | 2 | 1 |
|  | SACI059 | 0.125 | >8 |
|  | MbCI006 | 1 | 1 |
| Low virulence | UgCI223 | 2 | 1 |
|  | UgCI552 | 0.5 | 0.5 |
|  | SACI010 | 2 | ND |

ND: Not determined, isolate did not grow

**Supplemental Table 3.** Mice lung histology.

|  | Strain | Airways | Bronchial Epithelium/<br>Lamina Propria | Central<br>Alveoli | Inter-alveolar<br>septa | Peripheral<br>Alveoli | Granuloma |
| --- | --- | --- | --- | --- | --- | --- | --- |
| **High**<br>**virulence** | Non-infected | - | - | - | - | - | - |
| | KN99α | + | - | +++ | + | +++ | +++ |
|  | SACI012-1 | +/++ | - | ++ | - | ++ | +/++ |
|  | SACI012-2 | +/++ | - | ++ | - | + | ++ |
|  | SACI012-3 | ++ | - | ++ | - | +/++ | ++ |
|  | SACI012-4 | ++ | - | +++ | - | + | ++ |
|  | UgCI302-1 | + | + | +++ | - | ++/+++ | ++ |
|  | UgCI302-2 | + | - | +++ | - | +++ | +++ |
|  | UgCI387-1 | + | + | +++ | - | ++ | ++ |
|  | UgCI387-2 | ++ | - | +++ | - | ++ | ++ |
|  | UgCI395-1 | + | - | +++ | - | ++/+++ | ++ |
| **Intermediate**<br>**virulence** | UgCI538-1 | ++ | - | +++ | - | +++ | - |
|  | UgCI538-2 | - | - | ++ | - | + | -/+ |
|  | UgCI425-1 | ++ | +++ | ++ | ++ | ++ | ++ |
|  | UgCI425-2 | + | ++ | ++ | ++ | ++ | +++ |
|  | SACI059-1 | +/++ | ++ | ++ | ++ | ++ | +++ |
|  | MbCI006-1-1 | - | - | +/++ | - | + | ++ |
|  | MbCI006-1-1 | + | - | +/++ | - | + | ++ |
| **Low**<br>**virulence** | UgCI552-1 | - | - | - | - | - | - |
|  | UgCI552-2 | - | - | - | - | - | - |
|  | UgCI223-1 | + | ++ | + | ++ | + | +/++ |
|  | UgCI223-2 | - | - | + | - | + | + |
|  | SACI010-1 | - | - | + | - | -/+ | -/+ |
|  | SACI010-2 | - | - | + | - | + | -/+ |

Presence of *Cryptococcus*: - none, + scattered, ++ multi-focal, +++ confluent

**Supplemental Table 4.** Mice Brain Histology.

|  |  | LOCATION OF *CRYPTOCOCCUS* CELLS |  |  |  |  | IMMUNE RESPONSE |  |  |
| --- | --- | --- | --- | --- | --- | --- | --- | --- | --- |
|  | Strain | Meninges | Cerebrum | Cerebellum | Hippo/thalamus | Granuloma | PMNs | Macrophages<br>/Dendritic cells | Necrosis |
| **High virulence** | SACI012-1 | - | - | - | - | - | - | - | - |
|  | SACI012-4 | - | - | - | - | - | - | - | - |
|  | SACI012-2 | + | ++ | - | - | + | ++ | - | ++ |
|  | SACI012-3 | - | ++ | - | ++ | - | - | - | ++ |
|  | UgCI302-1 | + | - | ++ | - | - | + | + | - |
|  | UgCI302-2 | - | - | - | - | - | - | - | - |
|  | UgCI387-1 | - | - | - | - | - | - | - | - |
|  | UgCI387-2 | - | + | - | - | - | - | - | - |
|  | UgCI395-1 | - | - | - | - | - | - | - | - |
| **Intermediate virulence** | UgCI538-2 | - | - | - | - | - | + | - | - |
|  | UgCI538-1 | - | - | - | - | - | - | - | - |
|  | MbCI006-1 | - | - | - | - | - |  |  |  |
|  | MbCI006-2 | ++ | +++ | ++ | - |  | - | - | ++ |
|  | SACI059-1 | +++ | +++ | +++ | + | +++ | + | - | + |
|  | UgCI425-1 | +++ | ++ | + | + |  | - | - | - |
|  | UgCI425-2 | +++ | +++ | ++ | +++ | +++ | - | - | +++ |
| **Low virulence** | UgCI223-1 | +++ | ++ | ++ | ++ | - | - | - | +++ |
|  | UgCI223-2 | - | - | - | - | - | - | - | - |

Presence of *Cryptococcus*: - none, + scattered, ++ multifocal, +++ confluent
PMN: polymorphonuclear cells

**Supplemental Figure 1.**
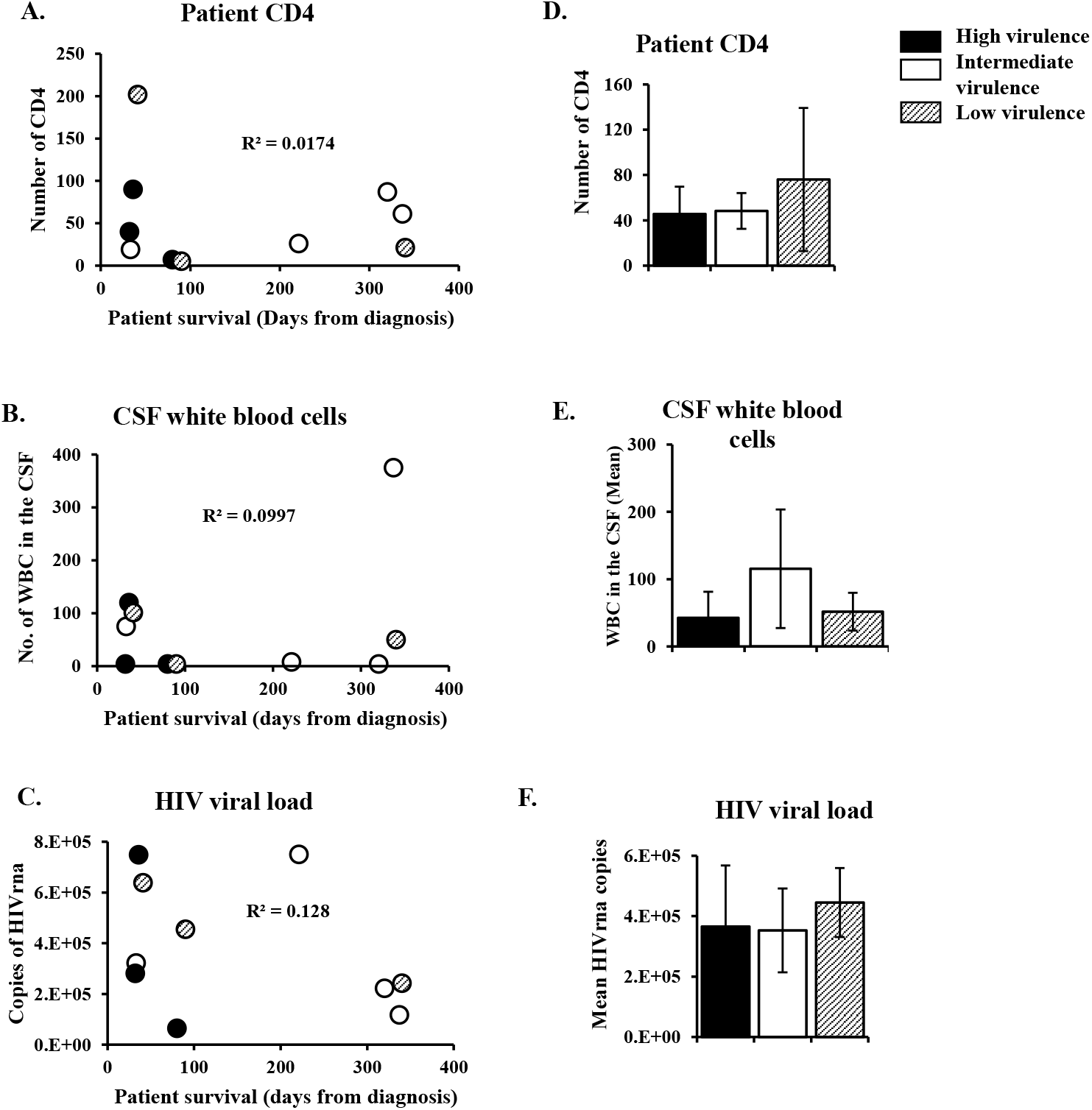
Human mortality is not determined by patient parameters. Clinical outcome was compared to baseline CM patient characteristics including CD4+ cells (A, D), white blood cell counts in the CSF (B, E), and HIV viral load (C, F). Graph A-C show individual patient parameters plotted against survival from diagnosis. Graphs D-F show patient parameters when classified by degree of virulence in the mouse model of cryptococcosis. Strains were classified in three groups: high (filled), intermediate (empty) and low virulence (striped) strains. Error bars represent standard error of the mean. CD4 cells: CD4+ T helper cells, CM: cryptococcal meningitis, CSF: cerebrospinal fluid, HIV: human immunodefiency virus, RNA: ribonucleic acid, WBC: white blood cells.

**Supplemental Figure 2.**
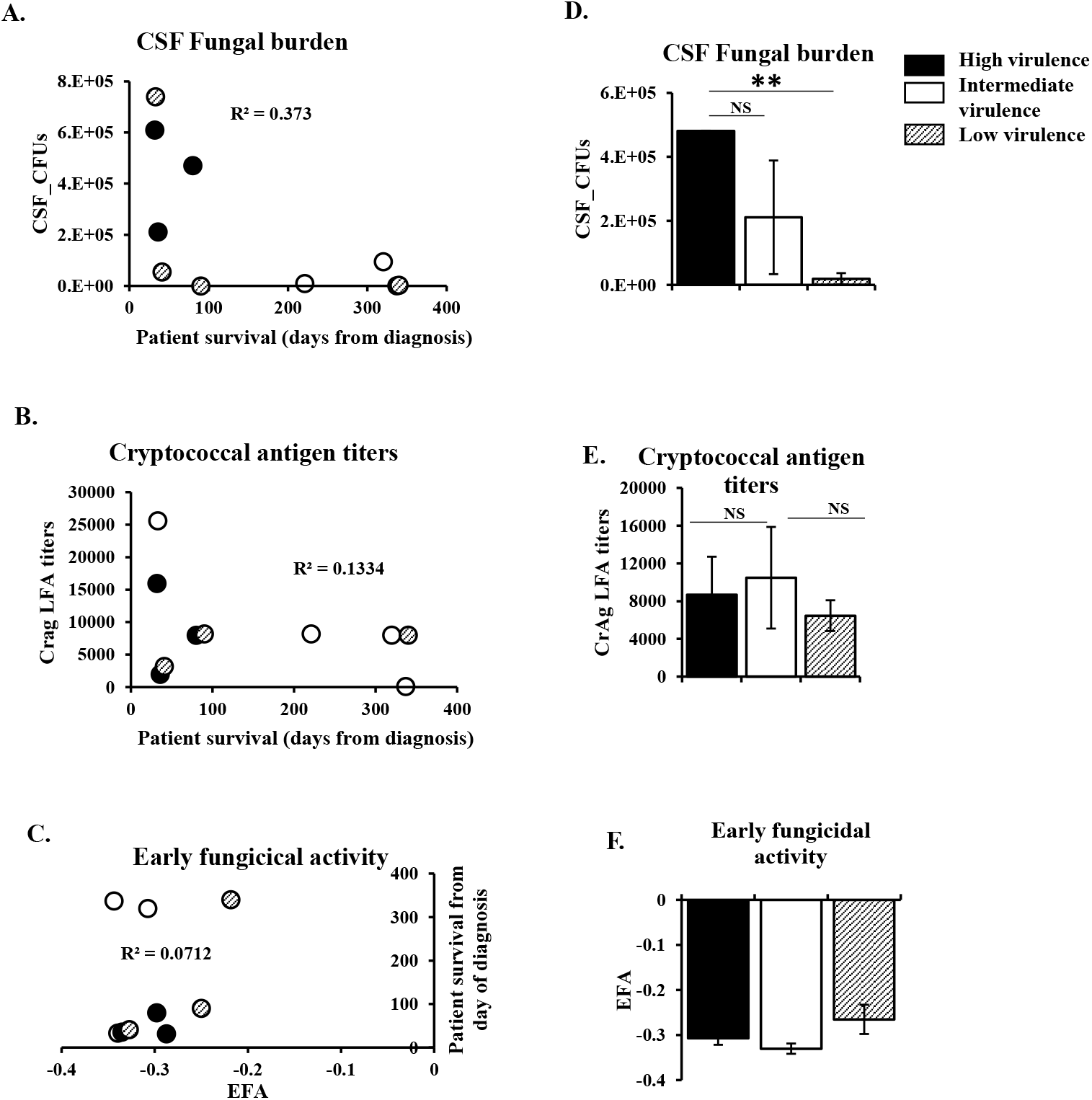
Human mortality is partly determined by fungal parameters. Clinical outcome was compared to fungal parameters including CSF-fungal burden determined by colony forming units (CFUs) (A, D), cryptococcal antigen titers in the CSF (B, E) and rate of fungal clearance determined by the early fungicidal activity (EFA) (C, F). Graphs A-C show individual patient parameters plotted against survival from diagnosis. Graphs D-F show patient parameters when classified by degree of virulence in the mouse model of cryptococcosis. Strains were classified in three groups: high (filled), intermediate (empty) and low virulence (striped). Error bars represent standard error of the mean. CFU: colony forming unit, CrAg LFA: cryptococcal antigen lateral flow assay, NS: not statistically significant. ** p < 0.01

**Supplemental Figure 3.**
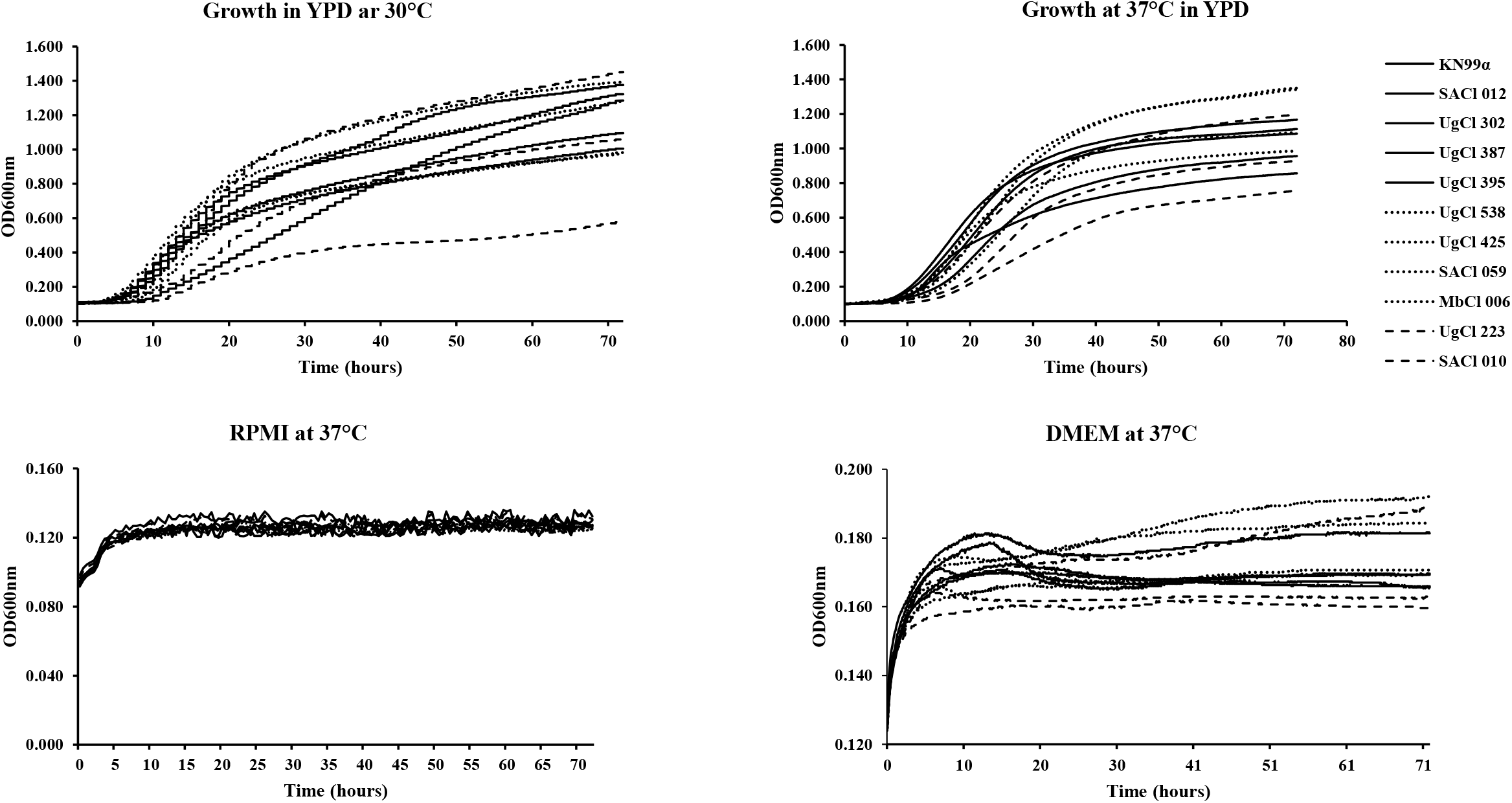
Absolute growth. Eleven *C. neoformans* clinical strains and the wild type control (KN99α) were grown overnight in YPD broth at 30°C, washed with PBS, and counted with a hemocytometer. 5 × 10^3^ yeast cells were transferred into 96-well plates containing 200 μl of either YPD (A, B), RPMI (C) or DMEM and grown at 30 or 37°C. Absorbances (OD600nm) were read during a 72 h incubation. Straight lines: high virulence strains, dotted lines: strains with intermediate virulence, dashed lines; low virulence strains. OD: optical density, DMEM: Dulbecco’s Modified Eagle’s Medium, RPMI: Roswell Park Memorial Institute medium, YPD: yeast extract-peptone-dextrose.

**Supplemental Figure 4.**
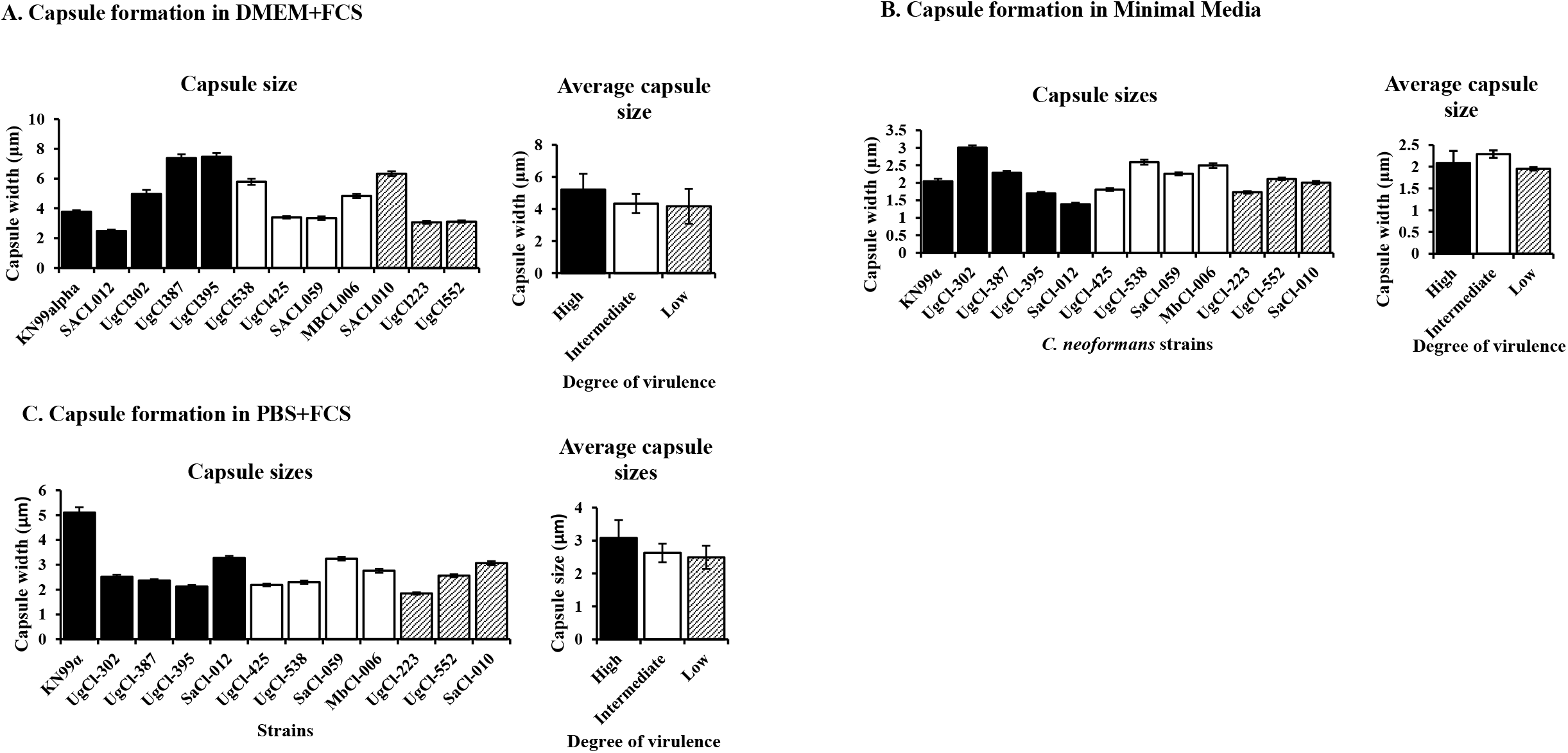
*In vitro* capsule formation. A *C. neoformans* wild type control strain and eleven clinical strains were induced to form capsule *in vitro* under three different conditions. **A)** Capsule induction in DMEM and serum: yeast cells were grown overnight in YPD, washed and then transferred into DMEM supplemented with FCS and incubated at 37°C, 5% CO_2_ for 5 days. **B)** Capsule induction in Minimal media: yeast cells were grown 22 h at 30°C in YPD, washed and resuspended in minimal media at a final concentration of 1 × 10^6^ cells/ml and incubated at 30°C with shaking (800rpm) in a thermomixer for 48 hours. C) Capsule induction in PBS and serum: yeast cells were grown overnight in YNB, washed and then transferred into PBS supplemented with FCS and incubated at 37°C, 5% CO2 for 48 hours. After incubation, yeast cells were fixed with 3.7% formaldehyde and observed under the microscope. The capsule width was defined as the difference between the diameter of the whole yeast cell (cell body and capsule) and the cell body diameter (no capsule) divided by 2. DMEM: Dulbecco’s Modified Eagle’s Medium, FCS: Fetal calf serum, PBS: Phosphate buffered saline, YNB: yeast nitrogen base medium, YPD: yeast extract-peptone-dextrose.

## References

1. Kambugu A, Meya DB, Rhein J, O’Brien M, Janoff EN, Ronald AR, Kamya MR, Mayanja-Kizza H, Sande MA, Bohjanen PR, Boulware DR. 2008. Outcome of cryptococcal meningitis in Uganda before and after the availability of HAART. Clin Infect Dis 46:1694–1701.

2. Rajasingham R, Smith RM, Park BJ, Jarvis JN, Govender NP, Chiller TM, Denning DW, Loyse A, Boulware DR. 2017. Global burden of disease of HIV-associated cryptococcal meningitis: an updated analysis. Lancet Infect Dis 17:873–881.

3. Perfect JR, Bicanic T. 2015. Cryptococcosis diagnosis and treatment: What do we know now. Fungal Genet Biol 78:49–54.

4. Bicanic T, Meintjes G, Wood R, Hayes M, Rebe K, Bekker LG, Harrison T. 2007. Fungal burden, early fungicidal activity, and outcome in cryptococcal meningitis in antiretroviral-naive or antiretroviral-experienced patients treated with amphotericin B or fluconazole. Clin Infect Dis 45:76–80.

5. Wiesner DL, Moskalenko O, Corcoran JM, McDonald T, Rolfes MA, Meya DB, Kajumbula H, Kambugu A, Bohjanen PR, Knight JF, Boulware DR, Nielsen K. 2012. Cryptococcal genotype influences immunologic response and human clinical outcome after meningitis. MBio e00196–12.

6. Aguiar P, Pedroso RDS, Borges AS, Moreira TA, Araujo LB, Roder D. 2017. The epidemiology of cryptococcosis and the characterization of *Cryptococcus neoformans* isolated in a Brazilian University Hospital. Rev Inst Med Trop Sao Paulo 59:e13.

7. Beale MA, Sabiiti W, Robertson EJ, Fuentes-Cabrejo KM, O’Hanlon SJ, Jarvis JN, Loyse A, Meintjes G, Harrison TS, May RC, Fisher MC, Bicanic T. 2015. Genotypic diversity is associated with clinical outcome and phenotype in cryptococcal meningitis across Southern Africa. PLoS Negl Trop Dis 9:e0003847.

8. Boulware DR, von Hohenberg M, Rolfes MA, Bahr NC, Rhein J, Akampurira A, Williams DA, Taseera K, Schutz C, McDonald T, Muzoora C, Meintjes G, Meya DB, Nielsen K, Huppler Hullsiek K. 2016. Human immune response varies by the degree of relative cryptococcal antigen shedding. Open Forum Infect Dis 3:ofv194.

9. Desnos-Ollivier M, Patel S, Raoux-Barbot D, Heitman J, Dromer F. 2015. Cryptococcosis serotypes impact outcome and provide evidence of *Cryptococcus neoformans* speciation. mBio 6:e003116.

10. Ferreira-Paim K, Andrade-Silva L, Fonseca FM, Ferreira TB, Mora DJ, Andrade-Silva J, Khan A, Dao A, Reis EC, Almeida MTG, Maltos A, Junior VR, Trilles L, Rickerts V, Chindamporn A, Sykes JE, Cogliati M, Nielsen K, Boekhout T, Fisher M, Kwon-Chung J, Engelthaler DM, Lazéra M, Meyer W, Silva-Vergara ML. 2017. MLST-based population genetic analysis in a global context reveals clonality amongst *Cryptococcus neoformans* var. *grubii* VNI Isolates from HIV patients in Southeastern Brazil. PLoS Neg Trop Dis 11:e0005223.

11. Day JN, Qihui S, Thanh LT, Trieu PH, Van AD, Thu NH, Chau TTH, Lan NPH, Chau NVV, Ashton PM, Thwaites GE, Boni MF, Wolbers M, Nagarajan N, Tan PBO, Baker S. 2017. Comparative genomics of *Cryptococcus neoformans* var. *grubii* associated with meningitis in HIV infected and uninfected patients in Vietnam. PLOS Neg Trop Dis 11:e0005628.

12. Boulware DR, Meya DB, Muzoora C, Rolfes MA, Huppler Hullsiek K, Musubire A, Taseera K, Nabeta HW, Schutz C, Williams DA, Rajasingham R, Rhein J, Thienemann F, Lo MW, Nielsen K, Bergemann TL, Kambugu A, Manabe YC, Janoff EN, Bohjanen PR, Meintjes G. 2014. Timing of antiretroviral therapy after diagnosis of cryptococcal meningitis. New Engl J Med 370:2487–2498.

13. Hommel B, Mukaremera L, Cordero RJB, Coelho C, Desjardins CA, Sturny-Leclère A, Janbon G, Perfect JR, Fraser JA, Casadevall A, Cuomo CA, Dromer F, Nielsen K, Alanio A. 2018. Titan cells formation in *Cryptococcus neoformans* is finely tuned by environmental conditions and modulated by positive and negative genetic regulators. PLoS Path 14:e1006982.

14. Dambuza IM, Drake T, Chapuis A, Zhou X, Correia J, Taylor-Smith L, LeGrave N, Rasmussen T, Fisher MC, Bicanic T, Harrison TS, Jaspars M, May RC, Brown GD, Yuecel R, MacCallum DM, Ballou ER. 2018. The *Cryptococcus neoformans* titan cell is an inducible and regulated morphotype underlying pathogenesis. PLoS Path 14:e1006978.

15. Smith KD, Achan B, Hullsiek KH, McDonald TR, Okagaki LH, Alhadab AA, Akampurira A, Rhein JR, Meya DB, Boulware DR, Nielsen K. 2015. Increased antifungal drug resistance in clinical isolates of *Cryptococcus neoformans* in Uganda. Antimicrob Agents Chemother 59:7197–204.

16. Nyazika TK, Hagen F, Machiridza T, Kutepa M, Masanganise F, Hendrickx M, Boekhout T, Magombei-Majinjiwa T, Siziba N, Chin, apos, ombe N, Mateveke K, Meis JF, Robertson VJ. 2016. *Cryptococcus neoformans* population diversity and clinical outcomes of HIV-associated cryptococcal meningitis patients in Zimbabwe. J Med Microbiol 65:1281–1288.

17. Haddow LJ, Colebunders R, Meintjes G, Lawn SD, Elliott JH, Manabe YC, Bohjanen PR, Sungkanuparph S, Easterbrook PJ, French MA, Boulware DR. 2010. Cryptococcal immune reconstitution inflammatory syndrome in HIV-1-infected individuals: proposed clinical case definitions. Lancet Infect Dis 10:791–802.

18. Okagaki LH, Strain AK, Nielsen JN, Charlier C, Baltes NJ, Chretien F, Heitman J, Dromer F, Nielsen K. 2010. Cryptococcal cell morphology affects host cell interactions and pathogenicity. PLoS Pathog 6:e1000953.

19. Zaragoza O, Garcia-Rodas R, Nosanchuk JD, Cuenca-Estrella M, Rodriguez-Tudela JL, Casadevall A. 2010. Fungal cell gigantism during mammalian infection. PLoS Pathog 6:e1000945.

20. Nichols CB, Perfect ZH, Alspaugh JA. 2007. A Ras1-Cdc24 signal transduction pathway mediates thermotolerance in the fungal pathogen *Cryptococcus neoformans*. Molecul Microbiol 63:1118–1130.

21. Kozel TR. 1995. Virulence factors of *Cryptococcus neoformans*. Trends Microbiol 3:295–9.

22. Alspaugh JA. 2015. Virulence mechanisms and *Cryptococcus neoformans* pathogenesis. Fungal Genet Biol 78:55–58.

23. Fernandes KE, Brockway A, Haverkamp M, Cuomo CA, van Ogtrop F, Perfect JR, Carter DA. 2018. Phenotypic variability correlates with clinical outcome in *Cryptococcus* isolates obtained from Botswanan HIV/AIDS patients. mBio 9:e02016–18.

24. Sionov E, Chang YC, Garraffo HM, Kwon-Chung KJ. 2009. Heteroresistance to fluconazole in *Cryptococcus neoformans* is intrinsic and associated with virulence. Antimicrob Agents Chemother 53:2804–15.

25. Robinson PA, Bauer M, Leal MAE, Evans SG, Holtom PD, Diamond DM, Leedom JM, Larsen RA. 1999. Early mycological treatment failure in AIDS-associated cryptococcal meningitis. Clin Infect Dis 28:82–92.

26. Bicanic T, Muzoora C, Brouwer AE, Meintjes G, Longley N, Taseera K, Rebe K, Loyse A, Jarvis J, Bekker LG, Wood R, Limmathurotsakul D, Chierakul W, Stepniewska K, White NJ, Jaffar S, Harrison TS. 2009. Independent association between rate of clearance of infection and clinical outcome of HIV-associated cryptococcal meningitis: analysis of a combined cohort of 262 patients. Clin Infect Dis 49:702–9.

27. Jackson A, Nussbaum J, Phulusa J, Namarika D, Chikasema M, Kenyemba C, Jarvis JN, Jaffar S, Hosseinipour MC, van der Horst C, Harrison TS. 2012. A phase II randomised controlled trial adding oral flucytosine to high dose fluconazole, with short-course amphotericin B, for cryptococcal meningitis. AIDS 26:1363–1370.

28. Jarvis JN, Bicanic T, Loyse A, Namarika D, Jackson A, Nussbaum JC, Longley N, Muzoora C, Phulusa J, Taseera K, Kanyembe C, Wilson D, Hosseiniour MC, Brouwer AE, Limmathurotsakul D, White N, van der Horst C, Wood R, Meintjes G, Bradley J, Jaffar S, Harrison T. 2014. Determinants of mortality in a combined cohort of 501 patients with HIV-associated cryptococcal meningitis: Implications for improving outcomes. Clin Infect Dis 58:736–745.

29. Bicanic T, Muzoora C, Brouwer AE, Meintjes G, Longley N, Taseera K, Rebe K, Loyse A, Jarvis J, Bekker L-G, Wood R, Limmathurotsakul D, Chierakul W, Stepniewska K, White NJ, Jaffar S, Harrison TS. 2009. Rate of clearance of infection is independently associated with clinical outcome in HIV-associated cryptococcal meningitis: analysis of a combined cohort of 262 patients. Clin Infect Dis 49:702–709.

30. Jarvis JN, Meintjes G, Rebe K, Williams GN, Bicanic T, Williams A, Schutz C, Bekker LG, Wood R, Harrison TS. 2012. Adjunctive interferon-gamma immunotherapy for the treatment of HIV-associated cryptococcal meningitis: a randomized controlled trial. AIDS 26:1105–13.

31. Chottanapund S, Singhasivanon P, Kaewkungwal J, Chamroonswasdi K, Manosuthi W. 2007. Survival time of HIV-infected patients with cryptococcal meningitis. J Med Assoc Thai 90:2104–11.

32. Sabiiti W, Robertson E, Beale MA, Johnston SA, Brouwer AE, Loyse A, Jarvis JN, Gilbert AS, Fisher MC, Harrison TS, May RC, Bicanic T. 2014. Efficient phagocytosis and laccase activity affect the outcome of HIV-associated cryptococcosis. J Clin Invest 124:2000–2008.

33. Zaragoza O, Chrisman CJ, Castelli MV, Frases S, Cuenca-Estrella M, Rodriguez-Tudela JL, Casadevall A. 2008. Capsule enlargement in *Cryptococcus neoformans* confers resistance to oxidative stress suggesting a mechanism for intracellular survival. Cell Microbiol 10:2043–57.

34. Yauch LE, Lam JS, Levitz SM. 2006. Direct inhibition of T-cell responses by the *Cryptococcus* capsular polysaccharide glucuronoxylomannan. PLoS Pathog 2:e120.

35. Rivera J, Feldmesser M, Cammer M, Casadevall A. 1998. Organ-dependent variation of capsule thickness in *Cryptococcus neoformans* during experimental murine infection. Infect Immun 66:5027–30.

36. Zaragoza O, Casadevall A. 2004. Experimental modulation of capsule size in *Cryptococcus neoformans*. Biol Proced Online 6:10–15.

37. Zaragoza O, Rodrigues ML, De Jesus M, Frases S, Dadachova E, Casadevall A. 2009. The capsule of the fungal pathogen *Cryptococcus neoformans*. Adv Appl Microbiol 68:133–216.

38. Clancy CJ, Nguyen MH, Alandoerffer R, Cheng S, Iczkowski K, Richardson M, Graybill JR. 2006. *Cryptococcus neoformans* var. *grubii* isolates recovered from persons with AIDS demonstrate a wide range of virulence during murine meningoencephalitis that correlates with the expression of certain virulence factors. Microbiol 152:2247–55.

39. Dykstra MA, Friedman L, Murphy JW. 1977. Capsule size of *Cryptococcus neoformans*: control and relationship to virulence. Infect Immun 16:129–135.

40. Littman ML, Tsubura E. 1959. Effect of degree of encapsulation upon virulence of *Cryptococcus neoformans*. Proc Soc Exp Biol Med 101:773–7.

41. Grossman NT, Casadevall A. 2017. Physiological differences in *Cryptococcus neoformans* strains *in vitro* versus *in vivo* and their effects on antifungal susceptibility. Antimicrob Agents Chemother 61.

42. Okagaki LH, Nielsen K. 2012. Titan cells confer protection from phagocytosis in *Cryptococcus neoformans* infections. Eukaryot Cell 11:820–6.

43. Litvintseva AP, Mitchell TG. 2009. Most environmental isolates of *Cryptococcus neoformans* var. *grubii* (Serotype A) are not lethal for mice. Infect Immun 77:3188–3195.

44. Desnos-Ollivier M, Patel S, Spaulding AR, Charlier C, Garcia-Hermoso D, Nielsen K, Dromer F. 2010. Mixed infections and in vivo evolution in the human fungal pathogen *Cryptococcus neoformans*. MBio 1:e00091–10.

45. Nielsen K, Cox GM, Wang P, Toffaletti DL, Perfect JR, Heitman J. 2003 Sexual cycle of *Cryptococcus neoformans* var. *grubii* and virulence of congenic a and α isolates. Infect Immun 71:4831–41.

